# Immune variation and host ontogeny shape virulence and transmission potential in a helminth parasite

**DOI:** 10.64898/2026.05.27.728195

**Authors:** Chloe A. Fouilloux, Heather Alexander, Patrick McNamara, Emma Polard, Ben Flanagan, Noelle Plantier, Dina Sokolovskaya, Gerrit Hoving, John Berini, Daniel I. Bolnick, Amanda K. Hund, Jessica L. Hite

## Abstract

Host immune strategies vary widely, yet how this variation shapes parasite virulence and transmission in natural systems remains unresolved. We tested whether host immune strategy governs macroparasite transmission timing and magnitude in *Schistocephalus solidus*, a cestode with a three-host life cycle. Over 13 months, we surveyed infection and immune activation (fibrosis) in second-intermediate host populations (threespine stickleback) from eight lakes. Host immune strategy (ranging from resistance to tolerance) strongly influenced whether and when parasites reached transmissible sizes, and shifted the synchrony between transmissible-stage peaks and definitive host availability. Across populations, transmission potential peaked at intermediate parasite burdens, consistent with virulence-transmission trade-off theory. These findings demonstrate that host immune phenotype alters epidemiological outcomes in complex life cycle parasites, by shifting both the magnitude and phenology of parasite growth to structure the effective transmission window.

## Introduction

Hosts and parasites are locked in a perpetual tug of war. To grow and transmit among hosts, parasites must successfully evade a wide array of host defenses (Ewald 1983; Antia et al. 1994). Hosts, in turn, must limit parasite induced damage without incurring excessive costs through immune activation or immunopathology (Graham et al. 2005; Viney and Cable 2011). Intuitively, to play this game, parasites should grow just enough to ensure transmission, while hosts should mount immune responses that effectively control infection that effectively control infection without excessive self-harm. Why, then, do some parasites cause severe disease or even kill their hosts? And why do some hosts mount seemingly maladaptive responses that lead to immunopathology? Answering these questions is essential not only for understanding host–parasite biology, but also for predicting how immune based interventions, including drugs and vaccines, shape parasite transmission and the evolution of virulence (Ewald 1983; Gandon et al. 2001; Dieckmann et al. 2002; Galvani 2003).

Evolutionary theory frequently leverages cost-benefit analyses to predict optimal phenotypes and to better understand why observed patterns deviate from these optima (Alizon et al. 2009; Lion and Metz 2018). While applications of this framework in immunology are gaining traction (Silva and Koella 2025), in evolutionary epidemiology the majority of this work has focused on parasite traits. For example, evolutionary theory predicts that virulence is an unavoidable consequence of selection to optimize transmission (Levin and Pimentel 1981; Anderson and May 1982; Bremermann and Pickering 1983; Ewald 1983; van Baalen and Sabelis 1995). Parasites must replicate (or grow) within their hosts and generate infective stages. However, higher replication intensifies exploitation of host resources, causes pathology, and can elicit stronger immune responses, all of which truncate the window over which transmission can occur. Parasites therefore face a trade-off: increased replication can elevate transmission rates, yet also imposes substantial costs. As a result, optimal parasite fitness occurs at intermediate levels of within host replication.

Numerous theoretical studies have focused on this trade-off hypothesis, especially in relation to public health decisions (Ewald 1983; Gandon et al. 2001; Dieckmann et al. 2002). Empirical tests, however, have produced mixed results (Alizon et al. 2009; Acevedo et al. 2019; Franz et al. 2025). Some studies have shown optimal transmission at an intermediate level of virulence (Jensen et al. 2006; de Roode et al. 2008) while others find positive relationships between within-host replication and virulence (Lipsitch and Moxon 1997) as well as between virulence and parasite transmission potential (Mackinnon and Read 1999; Messenger et al. 1999; Jensen et al. 2006). Such mixed empirical support has led some authors to question the generality of the virulence– transmission trade off (Ebert and Bull 2003; Koella and Turner 2007).

However, these inconsistencies may arise, in part, for multiple, potentially overlapping reasons. Parasites are among the most ubiquitous and diverse life forms on Earth (Windsor 1998). Yet, empirical tests of virulence trade-offs have sampled only a small subset of these systems, focusing primarily on directly transmitted microparasites (Antia et al. 1994; Levin 1996; Alizon et al. 2009; Acevedo et al. 2019). This focus makes sense given that the trade off theory was developed primarily for bacteria and viruses, with substantially less theoretical and empirical attention devoted to macroparasites, especially those with complex life cycles (Alizon et al. 2009; Viney and Cable 2011; Acevedo et al. 2019; Franz et al. 2025). These parasites differ fundamentally in how within host processes scale to transmission potential among host species (Box 1).

For macroparasites with complex life cycles, fitness is not optimized within a single host, but emerges through successful transmission across hosts (Woolhouse et al. 1995). To complete their life cycle, many of these parasites must undergo extended growth within a single host species, often lasting months to years, before reaching the stage capable of infecting the next host. Then, they must synchronize that development with discrete opportunities to encounter the next host species (Viney and Cable 2011). Additionally, for macroparasites, parasite size is often strongly correlated with egg production (Sinniah and Subramaniam 1991; Stear et al. 1995; Hollingsworth et al. 2015), increased disease severity (Schultz et al. 2006; Benesh 2023), and transmission (Tierney and Crompton 1992; Benesh and Hafer 2012; Froelick et al. 2021). Hence, selection on growth and development time is likely a strong factor regulating fitness in macroparasites.

Both empirical and theoretical work underscores that virulence and transmission are shaped by a complex interplay between parasite traits and host defense mechanisms (Van Baalen 1998; Gandon and Michalakis 2000; Gandon et al. 2001; Alizon et al. 2009). Immune variation is, however, rarely incorporated into empirical tests of the trade-off (Long et al. 2008; Alizon et al. 2009; Hite 2020). These gaps in knowledge are particularly acute for non-model organisms outside of controlled laboratory settings (Graham et al. 2011; Viney and Cable 2011; Duffy et al. 2021). A critical research endeavor is, therefore, to examine the complex interplay between immunity and parasites across a diverse array of host–parasite systems (Van Leeuwen et al. 2019).

Our goal here is to advance this endeavor by simultaneously quantifying relationships between immune variation, parasite burden (a proxy for virulence), and transmission potential in natural populations. As a case study, we use a tapeworm parasite (*Schistocephalus solidus*) with a complex life cycle involving copepods, threespine stickleback fish (*Gasterosteus aculeatus*), and piscivorous birds (Fig. 2). Threespine stickleback represent the only obligate host of the *S. solidus* infection sequence and thus a critical bottleneck for *S. solidus* transmission (Smyth 1946; Scharsack et al. 2007; Jakobsen et al. 2012). In its fish host, parasites must reach a minimum size (∼50 mg; Tierney and Crompton 1992; Barber and Svensson 2003) to become infective to definitive hosts, establishing a critical threshold for estimating *S. solidus* transmission potential.

The main barrier to reaching this threshold is host immunity, which varies within and among stickleback populations. Natural populations of stickleback exhibit heritable variation in immunity that ranges from resistance to tolerance (Weber et al. 2017b, 2022; Fuess et al. 2021; Hund et al. 2022). While tolerant populations permit parasite growth with limited immune activation, some populations have evolved a strikingly strong resistance phenotype (Weber et al. 2022). In resistant populations, hosts mount an intense fibrotic peritoneal response, characterized by granulomatous tissue encapsulating the tapeworm that effectively suppresses parasite growth. This fibrosis functions as an anti growth resistance mechanism, reducing parasite size by nearly two orders of magnitude relative to tolerant populations (Weber et al. 2017b; Hund et al. 2022). However, this potent immune defense is also associated with reduced nesting success in males and lower egg production in females (De Lisle and Bolnick 2021; Weber et al. 2022). Despite substantial costs to the host, peritoneal fibrosis is a common, evolutionarily conserved immune strategy (Vrtílek and Bolnick 2021).

Yet, despite extensive work with this well-studied host-parasite system, how these within host processes interact and scale up to shape population-level processes remains fragmentary (Heins and Baker 2011; Sprehn et al. 2015; Nordeide and Matos 2016). Beyond basic biology, a better understanding of these multi-scale interactions could help improve options to manage parasites with complex life cycles. Here, we build on previous work using short-term (13-month) time series focused on eight lakes to explicitly address three key questions:

1. How do the timing and magnitude of host immune response vary across infection status, host ontogeny, and season?
2. Does variation in immune responses affect synchrony between transmissible parasites and the abundance of definitive hosts (piscivorous birds)?
3. How do these immune mediated processes shape links between parasite burden (a proxy for virulence) and transmission *potential* to definitive hosts?

The multifaceted nature of the factors examined above makes *a priori* predictions for the first two questions challenging. However, for the third question we test the prediction from trade-off theory that transmission potential will peak at intermediate levels of parasite burden (biomass). In this system, parasite biomass represents a biologically meaningful metric for virulence because parasites commonly exceed 25% of host body mass (Arostegui et al. 2018) and greater parasite biomass is associated with increased exploitation of host resources and subsequent reductions in host growth, reproduction, and survival (Heins and Baker 2003; Bagamian et al. 2004).

While transmission *per se* remains challenging to quantify, infection prevalence can serve as a biologically reasonable proxy. Under common epidemiological assumptions, equilibrium prevalence approximates parasite fitness, often formalized as R_0_, the basic reproductive ratio (Anderson and May 1982). Thus, all else equal, infection prevalence can capture the same qualitative information as R_0_. Given this equivalence, we use infection prevalence when investigating links between virulence and transmission (see additional details in Box 1).

## Methods

### Natural history of the host-parasite system

*Schistocephalus solidus* (a diphyllobothrian cestode) is a trophically transmitted tapeworm with a three-host life cycle (Fig. 2D). Cued by light and temperature, free-swimming coracidia hatch from *S. solidus* eggs that must be ingested by their first-intermediate host, the cyclopoid copepod (Jakobsen et al. 2012). Infected copepods must be predated by a threespine stickleback (the only obligate host of the infection sequence), where the parasite rapidly burrows through the intestinal epithelium of the fish to enter the body cavity (Smyth 1946). In this second-intermediate host, *S. solidus* must reach a minimum size (∼50 mg, Tierney and Crompton 1992; Barber and Svensson 2003) to be infective to definitive hosts, which requires weeks to months (Barber and Svensson 2003; Scharsack et al. 2007). When piscivorous birds (e.g., loons) eat infected stickleback, mature parasites exit the fish and mate in the bird intestine, and eggs are released into the water in bird feces (Smyth 1946; Jakobsen et al. 2012; Fuess et al. 2021).

### Field survey

We sampled eight lakes near Bamfield, B.C. on Vancouver Island monthly over the course of one year, July 2023 – July 2024 (Fig. 1). These lakes were intentionally selected to capture variation in host immune responses across a resistance-tolerance gradient. At each monthly visit, we collected stickleback following standard methods (Marcogliese 1995; Weber et al. 2017a). Each month, we aimed to sample 50 adults and 50 juveniles per lake, although this target was occasionally limited during winter months (see <u>Supp.</u> Table 1 for monthly counts). Briefly, we deployed unbaited “Gee”-style minnow traps (¼″ mesh) along lake shorelines at 0.5–4.0 m depth and allowed them to soak overnight. When feasible, depending on lake conditions and season, we supplemented trapping with beach seining (¼″ mesh, 30′ × 6′ net) and hand-held dip nets (¼″ mesh). Once captured, stickleback were euthanized using MS-222 and immediately placed on ice until subsequent processing. We measured the standard length of each fish (SL; body length from the tip of the nose to the last vertebra) using digital calipers and weighed fish to the nearest 0.01 g. Smaller fish (typically < 20 mm SL) were preserved in ethanol and transported to the University of Connecticut for parallel investigations.

**Figure 1.**
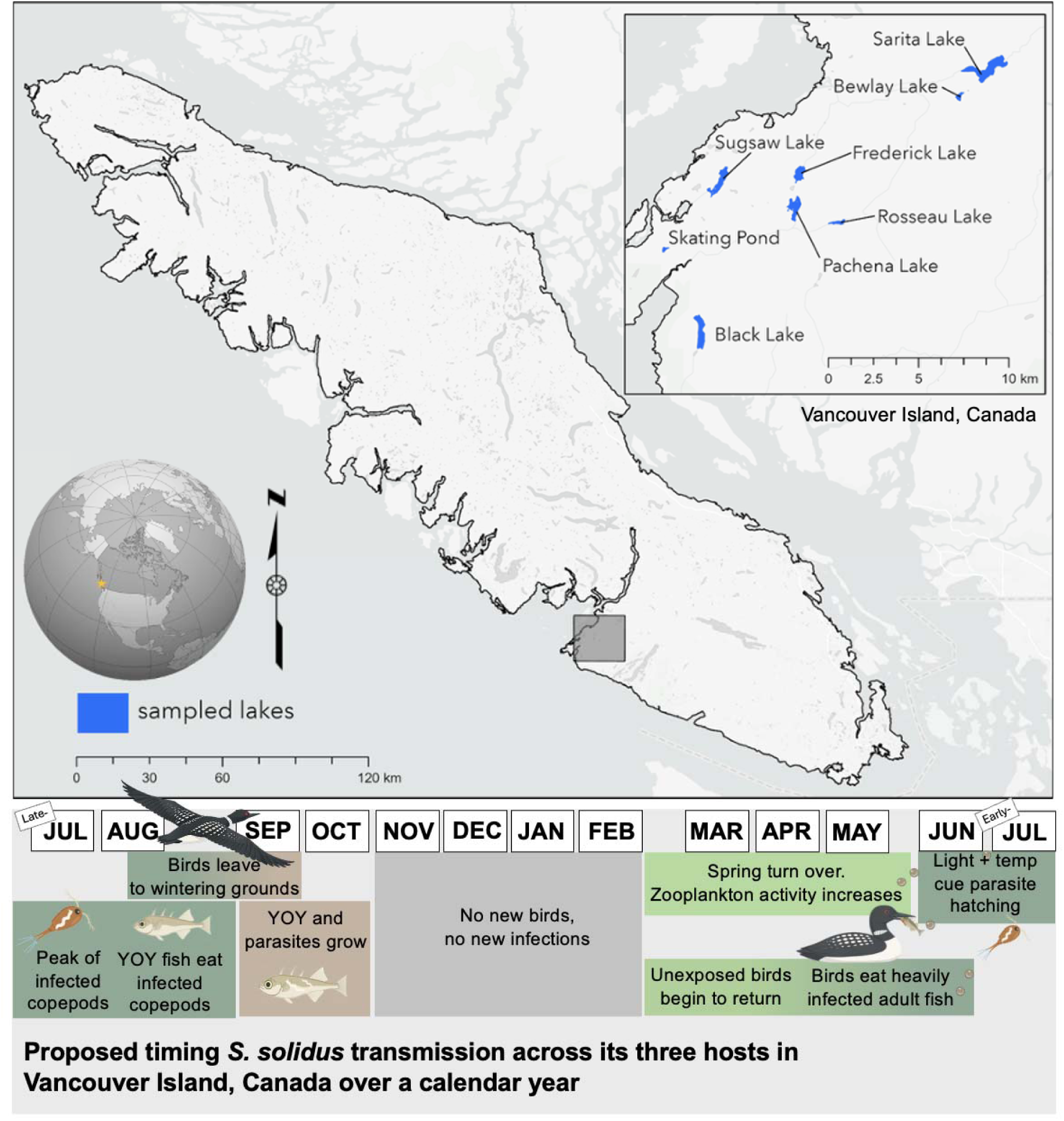
System overview. We surveyed stickleback populations from eight lakes near Bamfield, B.C. monthly across a calendar year (inset). The calendar depicts the proposed annual transmission cycle of *S. solidus* across its three hosts on Vancouver Island, anchored to the recruitment and infection history of a young-of-year stickleback.

**Figure 2.**
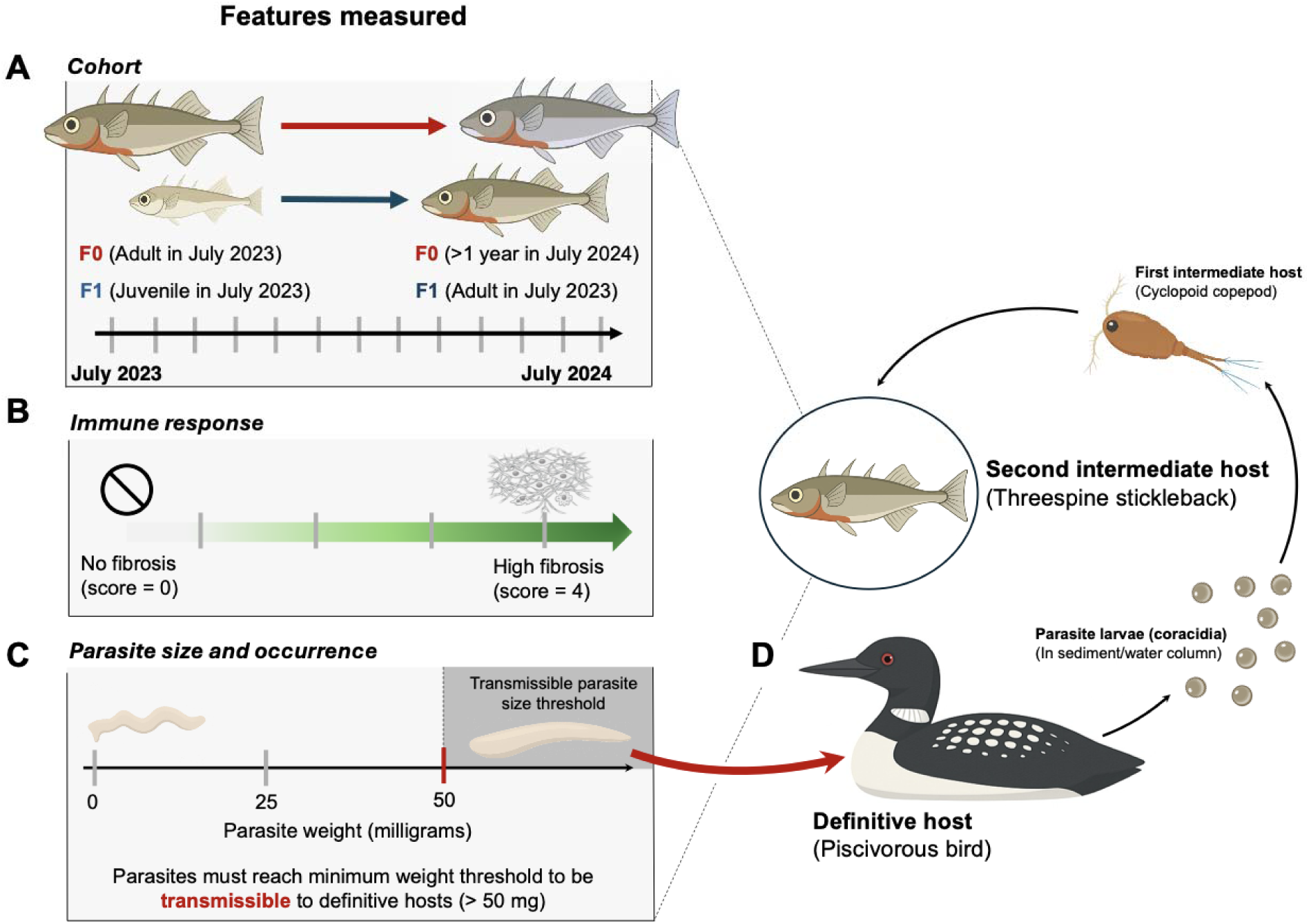
Study overview. The tapeworm, *Schistocephalus solidus,* is a trophically transmitted three-host parasite. In this study, we leverage (A) naturally-occurring stage-structured host populations of threespine stickleback where we examined (B) host immunity and (C) parasite infection dynamics across 13 months of sampling. (D) Transmission potential was assessed as the number of parasites exceeding the minimum transmissible size (>50 mg) in relation to definitive host availability across study lakes.

**Table 1.**
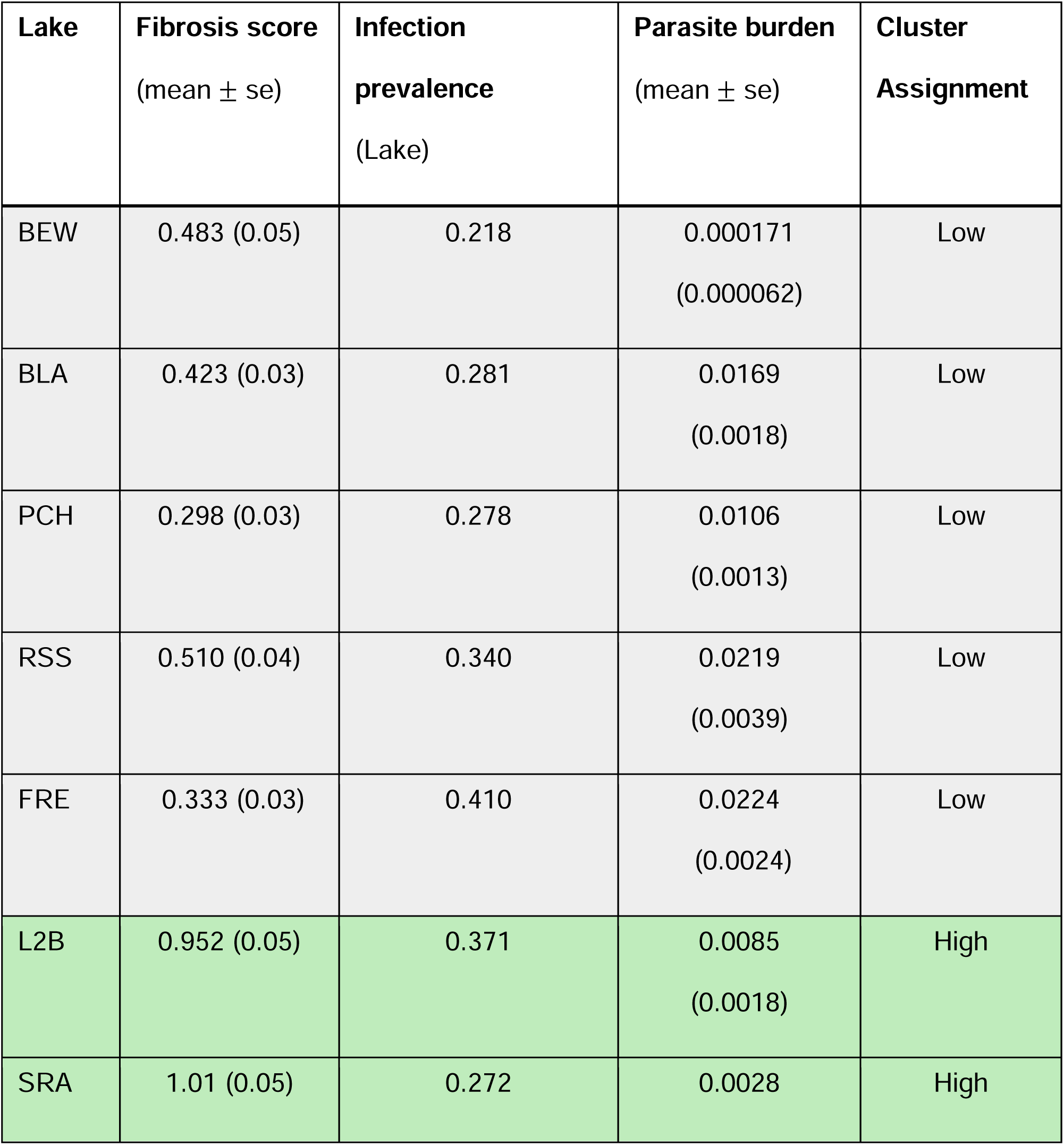
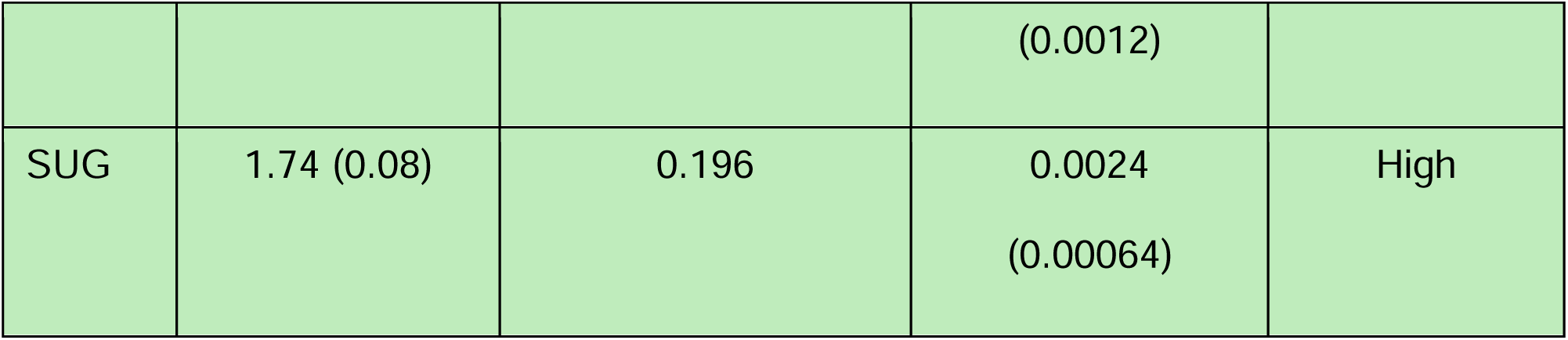
Lake clustering based on immune and infection dynamics.

### Host immune response

We visually assessed fibrosis and assigned a qualitative metric for fibrosis severity (ordinal 0-4) following previously established methods (Hund et al. 2022): 0 = no fibrosis; 1 = some fibrosis with limited organ movement; 2 = fibrosis, organs adhere together; 3 = fibrosis, organs adhered to each other and the peritoneal wall; 4 = severe fibrosis, making the cavity difficult to open (video reference: https://youtu.be/yKvcRVCSpWI; Hund et al. 2022). This fibrosis scoring system is repeatable across observers (r > 0.9; Bolnick et al. 2024) and correlates strongly with more labor-intensive laboratory measures of tissue stiffness (Flanagan, unpublished data). Because ethanol storage prevented reliable assessment of fibrosis, ethanol-stored individuals were excluded from analyses that used fibrosis as a response variable; thus, only fibrosis scores from dissected fish larger than ∼20 mm (range = 17.57-79.61 mm, n = 2470) were retained for this analysis, as preservation of smaller fish led to unreliable quantification.

### Infection metrics

We quantified infection prevalence (number infected fish/total fish surveyed) by visually diagnosing infection based on the presence of *S. solidus* tapeworms in the peritoneal cavity. To quantify parasite burden (total tapeworm mass per fish) for each individual host we dissected fish and weighed each individual tapeworm to the nearest 0.01 g. Parasites below this threshold were assigned a mass of 0.00005 g. For subsequent analyses, we used size-corrected parasite burden (total mass of tapeworms (g) divided by the total biomass of the host (g)) to account for population-level differences in host size as well as host growth over the course of the field survey.

### Definitive host sampling

To track spatio-temporal variation in the definitive hosts of *S. solidus*, we recorded bird calls for the duration of the sampling period (excluding July 2023 due to logistical constraints). Our goal was to estimate diversity, activity, and seasonality of piscivorous birds among lakes across the sampling period. In August 2023, we deployed autonomous recording units (ARUs) (Wildlife Acoustics™ Song Meter SM4 Acoustic Recorder) at each of the eight focal lakes. Each monitor was deployed as close to the lake shore as possible, at least 2 meters above the lake level, and oriented in an optimal direction to record bird songs on the lake and surrounding forest. The ARUs were scheduled to record for an on:off cycle of 30:30 minutes, and were programmed to record for the duration of the 12 months they were deployed (see Appendix 1 for pipeline details).

All sampling was conducted with IACUC approval (University of Connecticut) and Animal Care (BMSC) approval. Fish collections were approved by the Ministry of Forest, Lands, and Natural Resources Operations and Rural Development (Collections permit no. NA23-787881 and NA24-89569), Fisheries and Oceans Canada (License no. 139753), and the Huu-ay-aht First Nations (Heritage investigation permits no. 2023-013 and 2024-042). The sampling sites are located within the traditional territories of the Huu-ay-aht First Nations. Lakes include: Bewlay Lake (BEW), Black Lake (BLA), Frederick Lake (FRE), Pachena Lake (PCH), Skating Pond (L2B), Rosseau Lake (RSS), Sarita Lake (SRA), and Sugsaw Lake (SUG) (See Supp. Table 1 for GPS).

### Statistical analyses

Field sampling yielded direct measurements of host body size, fibrosis score, and parasite burden. Before statistical analyses, we derived two additional covariates from these measurements: population immune phenotype and individual cohort assignment. Because stickleback populations vary markedly in fibrosis expression and are inherently stage-structured, raw comparisons of infection dynamics across lakes and seasons are confounded by both factors. To resolve this, our analysis proceeded in two stages: we first characterized the host landscape by classifying lakes by immune phenotype using partitioning around medoids (PAM, Table 1) and assigning individual fish to generational cohorts (F0-F1-F2) using Gaussian mixture models (GMM). These classifications then served as covariates in three models examining how parasite burden, immune expression, and transmissibility vary across populations and through time (Fig. 2). All analyses were conducted in R version 4.5.1. Detailed statistical methods for all following sections can be found in Appendix 2.

*Q1. How do the timing and magnitude of host immune response vary across infection status, host ontogeny, and season*?

To examine variation in host immunity across host cohort and infection status across time, we used a generalized additive mixed model implemented in the `mgcv` package (v. 1.9.3, Wood 2026). Fibrosis score (0-4) was modeled as an ordered categorical response under the `ocat` family. Fixed effects included cohort (F0, F1) and the interaction between immune phenotype (high- vs low-fibrosis) and individual-level infection status in fish (binary, 0/1). Individual fibrosis scores measured at capture constituted the time-varying response variable, keeping immune phenotype and individual-level inference distinct. To capture nonlinear seasonal variation in fibrosis independently for high-vs. low-fibrosis populations, we included thin plate regression splines across months with separate smooth functions for each immune phenotype (k = 4). Lake identity was included as a random intercept through a regression-spline random-effect term (`s(Lake, bs = “re”)`).

To examine variation in parasite burden across host populations and through time, we used the stage structure and immune phenotype classifications outlined above and fit a Bayesian zero-inflated beta regression (`brms` v2.23.0, Bürkner 2021). Parasite burden across populations contained a high proportion of exact zeros, with 78.7% of sampled fish carrying no detectable parasites, a pattern common in naturally occurring parasite systems (Wilson et al. 2002; Blasco-Moreno et al. 2019). To account for this high proportion of structural zeros and to accommodate the bounded, continuous nature of the response variable (0, 1), we used a zero-inflated beta distribution.

*Q2. Does variation in immune responses affect synchrony between transmissible parasites and the abundance of definitive hosts (piscivorous birds)?*

We examined whether immune-mediated differences in transmissible parasite production align with peaks in presence of definitive hosts, linking immune dynamics to population-level transmission potential.

We quantified the total number of parasites available for trophic transmission to definitive hosts: we summed the number of transmissible-stage parasites (individual tapeworm mass ≥ 50 mg) across fish within each lake-month-cohort combination. Total number of fish with transmissible stage parasites were modeled using a negative binomial generalized additive mixed model (nb-GAMM in `mgcv`). Fixed effects included immune phenotype (high- vs. low-fibrosis) and host cohort (F0/F1). Immune-phenotype-specific thin-plate regression splines over sampling month were included to capture nonlinear seasonal variation independently for high- vs. low-fibrosis populations. Lake identity was included as a random intercept via `s(Lake, bs = “re”).` Models were fitted by REML.

To test whether transmissible parasite occurrence was synchronized with definitive host presence, we constructed monthly time series of transmissible parasite counts and piscivorous bird activity. For each month we summed the number of fish harboring transmissible parasites across lakes. We quantified definitive host presence as the number of piscivorous bird species detected acoustically and repeated the analysis using piscivorous call counts and loon call counts as alternative metrics. We restricted both series to the 12 months with paired acoustic data, excluding July 2023. Because bird recordings lacked lake-level resolution, we pooled both series across lakes and matched them by month.

We calculated monthly transmission potential as the product of the two time series and summed across months to obtain the observed test statistic. We generated a null distribution by permuting definitive host presence among months while holding the parasite series fixed. This decouples the series in time while preserving the observed values of each. We ran 9,999 permutations and calculated a standardized effect size as the difference between observed and mean null transmission potential divided by the null standard deviation. Because synchrony is directional, we report the one-tailed probability of obtaining a null value at least as large as observed.

*Q3. How do these immune mediated processes shape links between parasite burden (a proxy for virulence) and transmission potential to definitive hosts? –* We estimated links between virulence (parasite burden) and transmission *potential* to definitive hosts across host populations. To do so we integrated infection prevalence (proportion infected) across time using the area under the prevalence-time curve. This metric quantifies the size of epidemics that vary in length and shape over time (Van der Plank 1963, units: proportion x months). This index was strongly correlated with mean infection prevalence (Pearson’s r = 0.914, p = 0.0015; <u>Supp.</u> Fig. 2). We then examined links between integrated infection prevalence and virulence (parasite burden) across host populations (see Box 1 for further explanation).

We fit a quadratic regression of integrated infection prevalence on lake-level mean parasite burden of infected individuals. To test whether this relationship was unimodal rather than monotonic, we applied the Mitchell-Olds and Shaw test (MOStest in `vegan`, Oksanen et al. 2026).

## Results

### Immune responses are temporally coordinated across stage-structured populations

To determine whether hosts express immune responses dynamically across time (both seasonal and ontogenetic), we examined the strength of fibrosis expression across cohorts of high- and low-fibrosis populations across the calendar year. Overall, across our 13 months of sampling, 53% of the fish that we surveyed (1,315 out of 2,470) mounted a fibrotic immune response (fibrosis score > 0).

Population immune phenotype predicted individual fibrosis intensity: fish from low-fibrosis populations had significantly lower fibrosis scores than those from high-fibrosis populations, controlling for cohort, infection status, and month (Fig. 3A,B, GAMM, z = - 4.94, p < 0.001). Despite lower baseline fibrosis scores, infection in low-fibrosis populations was associated with a significant increase in fibrosis intensity (GAMM, interaction z = 4.35, p < 0.001), suggesting fibrosis is actively induced upon infection rather than constitutively expressed. When induced, fibrosis served as an effective anti-growth resistance mechanism: hosts that mounted a strong fibrosis response (score of 3-4; 10.8% of fibrotic individuals) never contained a transmissible tapeworm (<u>Supp.</u> Fig. 4).

**Figure 3.**
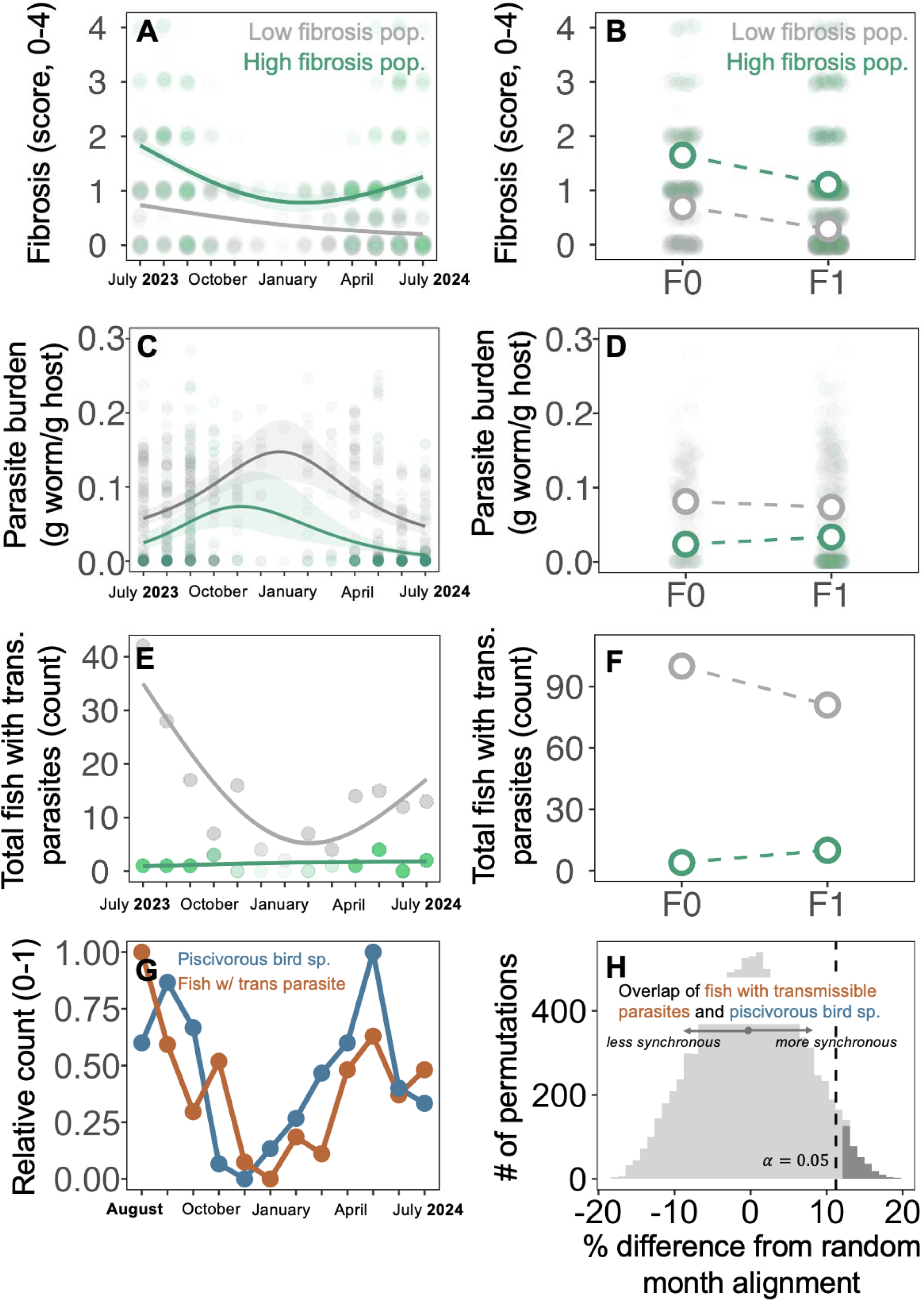
Immune phenotype and host ontogeny constrain pathogen virulence and transmission potential in a helminth parasite. (A, **B)** Fibrosis peaks in summer months (general additive mixed model (GAMM), edf = 2.67, p < 0.001) and is elevated in high-fibrosis populations and older cohorts (z = 5.44, p < 0.001). Fibrosis was scored in 2470 of 3397 fish. **(C, D)** Size-corrected parasite burden (g tapeworm/g host) peaks later and at higher magnitude in low-fibrosis populations (posterior estimate = 1.06, 95% CI 0.82 to 1.29; 2.6x higher), and F1 fish carry lighter burdens than F0 (posterior estimate = −0.22, 95% CI −0.39 to −0.04; ∼20% lower). Bayesian zero-inflated beta models were fit to all fish (n_Fish_ = 3397); estimates and smooths describe burden conditional on infection (n_Infected_ = 894). **(E, F)** The number of hosts with transmissible parasites peaks in summer months in low-fibrosis populations (nb-GAMM, edf = 2.12, p = 0.022); the cohort effect was not significant. Counts were aggregated by lake, month and cohort before fitting (n_Obs_ = 147, from 3397 fish across eight lakes). Smooths in (E) were fit separately by immune phenotype without cohort, for visualization only. **(G)** Monthly counts of fish harbouring transmissible parasites (>50 mg, orange) and of piscivorous bird species detected acoustically (blue), pooled across all eight lakes and normalized so that the lowest month equals zero and the highest equals one. **(H)** Observed transmission *potential* (overlap between piscivorous bird species count and fish with transmissible parasites) exceeded random temporal alignment by 12.1% (SES = 1.82, one-tailed p = 0.035). Histogram shows 9,999 permutations in which piscivorous species counts were reshuffled among months with the parasite series held fixed. Dashed line, α = 0.05 significance threshold; dark shading, permutations at or above the observed value. Green, high-fibrosis populations; grey, low-fibrosis populations (A–F). Transparent points, raw individual-level data (A–D) or aggregated lake × month × cohort counts (E). Open circles, cohort means ± SE, with SE masked by point size (B, D, F). Smooths fit with a general additive model (A, E) or Bayesian posterior prediction (C).

**Figure 4.**
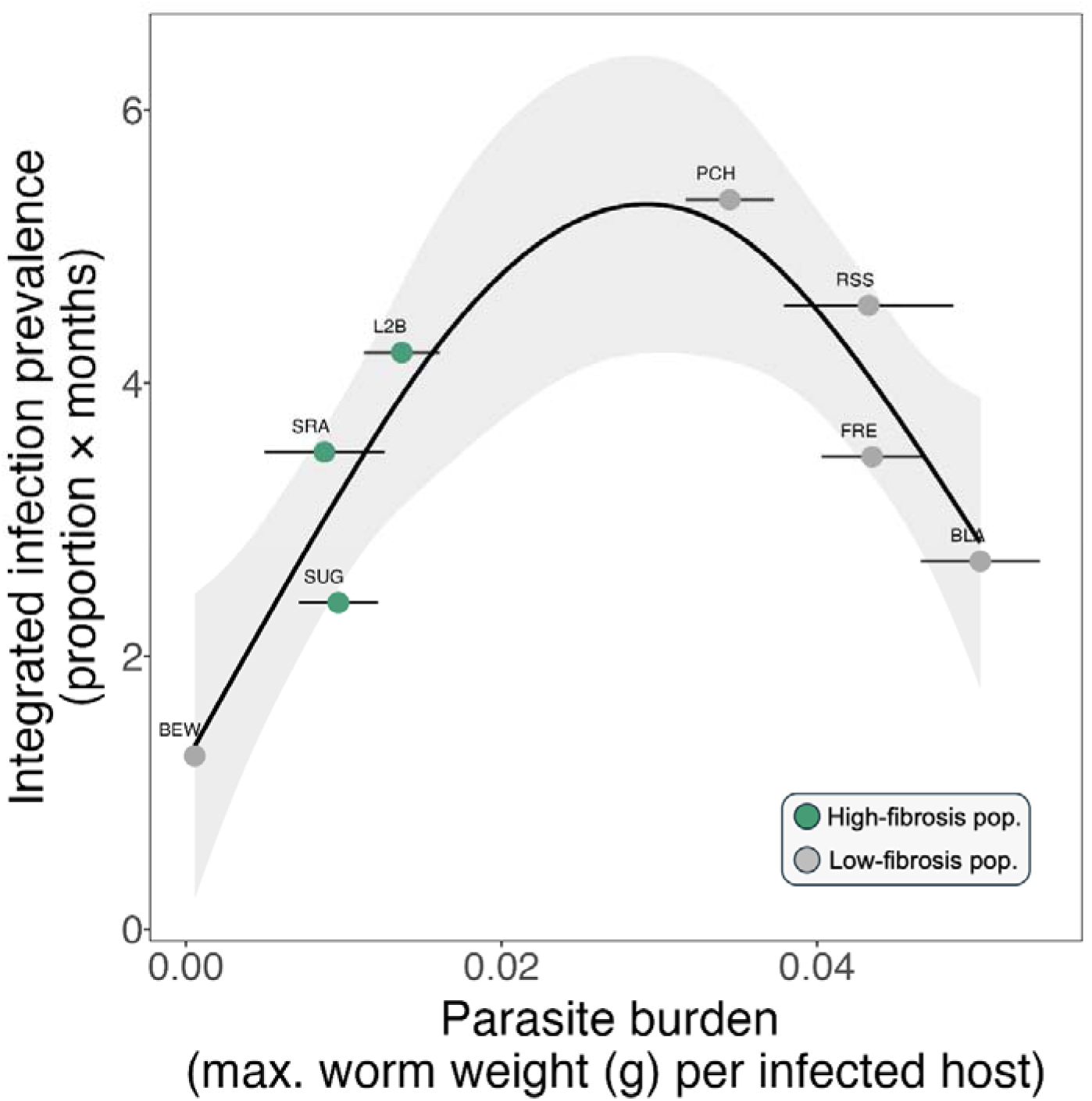
Transmission potential peaks at intermediate levels of parasite burden, a metric for virulence. The observed unimodal relationship between integrated infection prevalence (a proxy for transmission potential) and parasite burden is consistent with theoretical expectations under a virulence-transmission trade-off (MOS test, combined p = 0.013; see Box 1). For virulence, we use the maximum parasite burden, size-corrected for variation in host biomass (g tapeworm/g host), averaged over the time series for each lake. As a proxy for transmission potential to definitive hosts, we use integrated infection prevalence (proportion × days) calculated as the area under the prevalence-time curve. Data span July 2023 to July 2024 across 8 lakes on Vancouver Island. Point ranges (mean ± SE) are colored by immune phenotype (Green, high-fibrosis populations; grey, low-fibrosis populations). Smooth fit with a generalized additive model for visualization purposes only.

Across 13 months of sampling, < 1% of fish (n_HighFib_= 14/1420) from high-fibrosis populations harbored transmissible parasites compared to ∼10% from low-fibrosis populations (n_LowFib_ = 181/1977). This order of magnitude difference substantiates that fibrosis significantly (nb-GAMM, z = 4.873, p < 0.001) limits the number of parasites capable of onward transmission (Fig. 3E; Weber et al. 2022). There is clear temporal structure in fibrosis, with high- and low-fibrosis populations exhibiting mid-summer peaks across all eight lakes (Fig. 3A, July-August, GAMM, edf ≈ 2.67, p < 0.001; <u>Supp.</u> Fig. 3).

In addition to these seasonal dynamics, fibrosis also varied strongly across host age classes. Older cohorts (F0) across populations exhibited significantly higher fibrosis scores than young-of-year fish (Fig. 3B, GAMM, z = 5.44, p < 0.001). Nevertheless, our data reveal that young juveniles are capable of mounting strong fibrotic responses early in ontogeny (e.g., fibrosis scores ≥3 observed in July and August; the smallest confirmed fibrotic individual measuring 19.78 mm).

### Parasite burdens are higher and peak later in low-fibrosis populations

In total, we found 1,759 individual tapeworms across 3,397 sampled fish across our 13 months of sampling. Across lakes, mean *S. solidus* infection prevalence was 27.3% (range = 18.1 – 36.4%) and parasite infection intensity (number of tapeworms/infected host) ranged from 1–30 parasites (mean = 1.9 tapeworms ± 0.063 SE). The biomass of plerocercoid tapeworms dissected from stickleback ranged from 0.00005 g to 0.33 g (mean = 0.028 g ± 0.0013 SE). Both high- and low-fibrosis populations showed unimodal trajectories of parasite burden across time, peaking broadly in winter months. However, populations differed in the onset and amplitude of this peak: burden peaked later in low-fibrosis populations (Fig. 3C, peak: Month 6.75, ∼late December/early January) than in high-fibrosis populations (peak: Month 5, ∼November), and infected individuals from low-fibrosis populations carried substantially higher burdens (Fig. 3D, posterior estimate = 1.06, 95% CI = 0.82 to 1.29; 2.6x higher).

Across populations, younger cohorts (F1) tended to carry lighter burdens than older cohorts (F0) among infected individuals (Fig. 3D, posterior estimate = −0.22, 95% CI = −0.39 to −0.04; ∼20% lower burden in F1). We detected tapeworms in young-of-year (F1) fish as small as 12.66 mm in July, indicating that infection occurs early in stickleback development (similar to patterns in Heins et al. 2011, Wohlleben et al. 2022).

### Transmissible parasites occur most frequently in summer months, coinciding with peak definitive host activity

The timing of parasite transmission in natural systems depends on the temporal overlap between infective parasites in intermediate hosts and the occurrence of definitive hosts. We quantified seasonal variation in the frequency of fish carrying transmissible tapeworms and compared these patterns to acoustic occupancy data of piscivorous birds across BAMF-area lakes.

The number of fish carrying transmissible tapeworms varied significantly across months in low-fibrosis populations, peaking in summer (Fig. 3E, nb-GAMM, edf = 2.12, p = 0.022). These peaks coincided with periods of elevated piscivorous bird presence (measured here as piscivorous species counts, Fig. 3G). Observed overlap between fish with transmissible parasites and piscivorous bird species exceeded the month-permuted null (Fig. 3H, observed = 2005, null mean = 1788, SES = 1.82, one-tailed p = 0.035), 12.1% higher than expected under random temporal alignment. Call-based measures of definitive host presence gave the same direction with weaker support (piscivore calls: SES = 1.36, p = 0.097; loon calls: SES = 1.16, p = 0.137).

### Transmission *potential* to definitive hosts is optimized at intermediate parasite burdens

Across host populations, we found that intermediate levels of parasite burden (virulence; maximum tapeworm biomass (g) divided by the biomass of the host (g)) have the highest transmission potential (integrated infection prevalence, Fig. 4) (see Box 1 for details). The relationship was unimodal, with a peak at 0.029 g (MOS test, combined p = 0.013; F = 26.50, p = 0.004 at the lower bound, F = 16.68, p = 0.010 at the upper bound). Both boundary tests were rejected, confirming an interior optimum rather than a monotonic relationship with incidental curvature.

## Discussion

Understanding natural variation in host immune strategies and parasite traits such as virulence and transmission remains a central challenge in epidemiology, particularly for parasites with complex life cycles. Our study demonstrates that host ontogeny and immune strategies jointly shape parasite transmission potential in a naturally occurring macroparasite system. Population level differences in immune expression can impose strong constraints on parasite growth and development by altering the timing and mode of host defense. These immune mediated constraints restrict ability of parasites to reach infective thresholds. Together, these within-host processes scale up to shape population-level processes such that transmission potential to definitive hosts is maximized at intermediate infection burdens.

The timing and magnitude of host immune response varied across infection status, host ontogeny, and season. The temporal resolution of this study provides new insight into an evolutionarily conserved immune response, fibrosis (Vrtílek and Bolnick 2021). Across all populations, mean fibrosis peaked in summer months (Fig. 3A, Supp. Fig. 3). This temporal pattern reflects both parasite exposure and host life history. We found that young-of-year stickleback (F1s) are capable of expressing fibrosis, with the smallest confirmed fibrotic individual measuring 19.78 mm. The fibrotic response in juveniles is particularly notable given the substantial costs associated with immune responses in general (Zuk and Stoehr 2002; Schmid-Hempel 2003; Sadd and Schmid-Hempel 2009; Parker et al. 2011) and fibrosis in particular (De Lisle and Bolnick 2021; Weber et al. 2022; Matthews et al. 2023). Overall, however, older cohorts mounted significantly stronger fibrotic responses than young-of-year fish.

Theoretically, this is counterintuitive, as the costs associated with infection prior to reproduction are more damaging than those post-reproduction. Thus, selection should favor strong immune responses in juveniles (Ashby and Bruns 2018). Yet, across a diverse array of taxa, juveniles are typically more susceptible to parasites (Armitage and Boomsma 2010; Garbutt et al. 2014; Ashby and Bruns 2018). Our data suggest that this apparent data-theory disconnect may arise because stage-specific susceptibility and immune responses are subject to strong selection on timing: immune investment should peak when exposure risk or parasite growth potential is highest (Lochmiller and Deerenberg 2000; Schmid-Hempel 2003; Alizon 2008). Such temporal variation can arise through several factors and is likely pronounced in most host-parasite systems, but remains difficult to capture with standard mathematical models and empirical studies (Cornell et al. 2008).

Our study helps address these gaps and link temporal variation in the timing of immune responses to variation in parasite growth and development. For both low and high fibrosis populations, we found a unimodal pattern of parasite growth across time. However, the timing and amplitude were significantly different. In low-fibrosis populations, the absence of a strong fibrotic brake allowed parasites to capitalize on the full growth window. Low fibrosis (tolerant) stickleback populations sustained approximately threefold higher parasite burdens, with peak parasite growth occurring roughly two months later than in high fibrosis populations. Parasite burdens increased steadily through autumn and into December-January (Fig. 3A). The subsequent decline in burden mid-winter may reflect selective removal of heavily infected hosts through infection-associated mortality (Heins et al. 2011) or predation (by both non-transmitting predators and piscivorous birds) (Fig. 3A, grey line).

In contrast, the pattern of parasite development in high-fibrosis populations was right-skewed and compressed (Fig. 3C). This skew may arise from the rapid activation of fibrosis, which can begin within 24 hours of *S. solidus* exposure (Hund et al. 2022), creating an early developmental brake that limits parasite growth. Thus, shape of the unimodal distribution in resistant populations reveals how parasites stunted by fibrosis never recovered sufficiently to reach transmissible sizes (fewer than 1% of resistant fish harbored transmissible parasites; see also Weber et al. 2022). In these populations, the subsequent decline in parasite burden likely reflects the growth of hosts across time (increasing the denominator as the host grows and the parasite remains unchanged). Together, these dynamics describe two qualitatively distinct trajectories: one in which parasites exploit the full seasonal window before host costs terminate the association (low-fibrosis populations) and one in which the immune response arrests parasite development from the outset (high-fibrosis populations).

Did variation in these immune responses shift synchrony between transmissible parasites and the abundance of definitive hosts (piscivorous birds)? To reach a transmissible stage, *S. solidus* must survive immune-mediated growth suppression and attain a sufficient size to be transmissible (>50 mg; Tierney and Crompton 1992). Yet, larger *S. solidus* impose greater costs on fish (Schultz et al. 2006; Heins and Baker 2011) and winter ice cover can eliminate opportunities for transmission at northern latitudes (Nordeide and Matos 2016). Thus, parasites face a narrow window in which they must reach transmission-ready size before either host mortality or seasonal constraints on transmission terminate the association.

Across 13 months, we found transmission-ready tapeworms were significantly more frequent in the summer (Fig. 3E), aligning with previous work examining *S. solidus* growth in stickleback on monthly scales (Baer et al. 2022; Fig. 3E). When framed against piscivorous bird occurrence (species richness, call counts), transmission-ready stages occur in synchrony with definitive hosts (Fig. 3G, H). The timing of this peak to overlap with the occurrence of piscivorous birds is crucial, as *S. solidus* must reach its definitive host to sexually reproduce.

Our results highlight the challenges (and opportunities) of linking ecological factors with transmission across the entire life cycle. Even with detailed data from the bird survey, quantifying the abundance of piscivorous birds remained challenging. These results join a small set of studies documenting temporal overlap between sequential hosts in trophically transmitted parasites. For example, Amundsen et al. (2003) found peak infection of Arctic charr by the cestode *Cyathocephalus truncatus* coincided with a host diet shift toward amphipods, the parasite’s intermediate host. In trematodes, Nikolaev et al. (2020) found metacercarial infection of mussels peaked with seabird autumn migration. Our system differs in an important respect: the timing of transmission-ready stages is set not only by parasite development but by the seasonal immune phenology of the intermediate host. Temporal heterogeneity in infection risk arises from both ecological drivers of exposure and biological variation in host susceptibility, which often vary seasonally and across development (Zuk and Stoehr 2002; Cornell et al. 2008; Downie et al. 2021). Despite this, few studies have simultaneously tracked immune expression and parasite growth through time in wild populations (but see Cornell et al. 2008).

Together, our results highlight links between immune variation, parasite burden (a proxy for virulence) and transmission potential to definitive hosts. While theory predicts that parasite fitness should be maximized at intermediate levels of exploitation, empirical support for virulence-transmission trade-offs has been mixed, particularly for macroparasites with complex life cycles. Our findings suggest that host immune strategies are an important and often overlooked component of this relationship. By altering the timing and extent of parasite growth, immune-mediated constraints shape whether parasites reach the infective stage and, in turn, how within-host processes scale up to transmission opportunities among hosts.

Of course, we fully acknowledge that the sample size is limited and these metrics necessarily abstract over biological dimensions. Direct estimates of both virulence and transmission are notoriously difficult to obtain in natural populations, particularly for parasites with complex life cycles. Our goal was therefore not to quantify every process contributing to these parameters, but rather to examine population-level variation in biologically informed proxies that can be compared across lakes and seasons. We used parasite burden as a proxy for virulence based on extensive laboratory evidence linking parasite biomass to host harm (Heins and Baker 2003; Bagamian et al. 2004).

We also used integrated prevalence of infective parasites in stickleback as a biologically informed proxy for transmission potential. In parasites with complex life cycles, infection prevalence in second intermediate hosts does not directly quantify transmission to definitive hosts. And, logistically, we cannot directly measure establishment, reproduction, or life-cycle completion within the definitive hosts. Instead, infection prevalence (integrated over the time series of the field survey) captures the cumulative outcome of multiple processes, including exposure from copepods, parasite growth and persistence within stickleback, host mortality, recruitment, and parasite-mediated predation. However, theory suggests that infection prevalence can provide a biologically informed estimate of transmission potential (see detailed explanation in Box 1) and enables comparisons across populations and seasons (Anderson and May 1982; Alizon et al. 2009).

Even with these limitations, our findings provide a rare glimpse into links between host immunity and key parasite traits in a natural macroparasite system. This approach demonstrates that biologically informed measures of parasite burden and transmission potential can reveal how within-host processes scale up to shape population-level transmission dynamics in natural host-parasite systems.

### Limitations and future directions

As a field survey, this study necessarily integrates multiple ecological processes that cannot be separated mechanistically. First, we focused on temporal variation in host immunity, parasite growth, and infection dynamics rather than identifying the specific environmental drivers underlying those patterns. Our ongoing work focuses on understanding the potential environmental correlates linked with host and parasite traits (Hund et al., unpublished data).

Additionally, due to logistical constraints, individual fish were not marked, so monthly lake surveys represent population averages. While this gives us a rich basis to understand differences in parasite growth and immune activation from a population-level perspective, we did not track individual growth based on mark-recapture data. We were able to address this limitation, in part, by leveraging dynamic energy budget growth curves (Wright et al. 2004) for stickleback, allowing us to infer cohort identity (F0-F1-F2). However, future work may include mark-recapture methods to track the growth of stickleback individuals across time. This can be of value, as tapeworms are known to modify their life-history strategy according to host age (Viney and Cable 2011). For example, the helminth *Polystoma integerrimum* can plastically adjust its development window from three weeks to three years depending on the stage of the tadpole host it infects (Gallien 1935). How (and if) *S. solidus* regulates its growth as a function of host age upon infection is unknown, but a fascinating avenue for future work.

## Conclusion

While parasitologists have understood that parasites with complex life cycles must navigate multiple hosts to reach maturity, the factors that mediate transmission across space and time remain poorly resolved (Nordeide and Matos 2016). Together, this work highlights the value of natural history data in a well-studied system as a case study to help address critical gaps in the understanding of host-parasite systems. As anthropogenic disturbances increasingly alter the ecology of host-parasite interactions (Poulin et al. 2026), such insight has become essential for managing disease risk in natural populations (Plowright et al. 2024). Moreover, for helminths that affect wildlife, livestock, and humans alike, clarifying how ecological processes mediate virulence-transmission relationships is necessary to evaluate whether interventions can constrain parasite growth and transmission without favoring increased harm to hosts (Gandon et al. 2001; Hite 2020).

## Acknowledgements

We thank Sebastian Schrieber, Anieke Van Leeuwen, Kayla King, and Clay Cressler for helpful conversations; National Science Foundation EEID (No. 2243076 to AKH, DIB, JLH).

## Statement of authorship

CAF performed all statistical analyses and visualizations and prepared the first draft of the manuscript, in collaboration with JLH. HA, DB, JBIII. HA, DB, JBIII, and JLH led field work, dissections, and data management. PM, EM, NP, and DS contributed substantially to field sampling, dissections, and data entry. JBIII designed and led piscivorous bird monitoring and analysis, with substantial contributions from GH. All authors contributed critically to manuscript revisions and approved the final submission.

## Data accessibility

All data and code are publicly available on EDI (Fouilloux et al. 2026, DOI: 10.6073/pasta/a75d5c422a5c67dacd1c6222ba5f36ce)

## Box 1. Examining Virulence-Transmission Trade-offs in Macroparasite Systems

Can infection prevalence serve as a proxy for parasite fitness? Here, we illustrate how, under common epidemiological assumptions, equilibrium prevalence approximates parasite fitness, often formalized as, the basic reproductive ratio. This equivalence motivates the use of infection prevalence when investigating links between virulence and transmission **(Fig. 4).**

Both transmission rates and R_o_ are notoriously difficult to quantify, especially for parasites with complex life cycles that must infect multiple host species sequentially. For trophically transmitted parasites, no single virulence optimum emerges across taxonomically distinct hosts (Woolhouse et al. 1995). However, in the case study used here, the intermediate hosts, stickleback fish, represent the final bottleneck before parasites can realize fitness gains in their definitive avian hosts (Wedekind 1997). To reach this stage, parasites must both survive immune-mediated growth suppression and attain a sufficient size to be transmissible (>0.05 g; Tierney and Crompton 1992). Although we can not quantify transmission directly, we can quantify parasite burden within hosts, the number of parasites that reach transmissible stages, and the prevalence of infected (intermediate) hosts.

For a broad class of epidemiological models with endemic infection, equilibrium prevalence is a monotonic function of parasite fitness, such that parasites with higher ft_0_ generate larger outbreaks and higher infection prevalence.

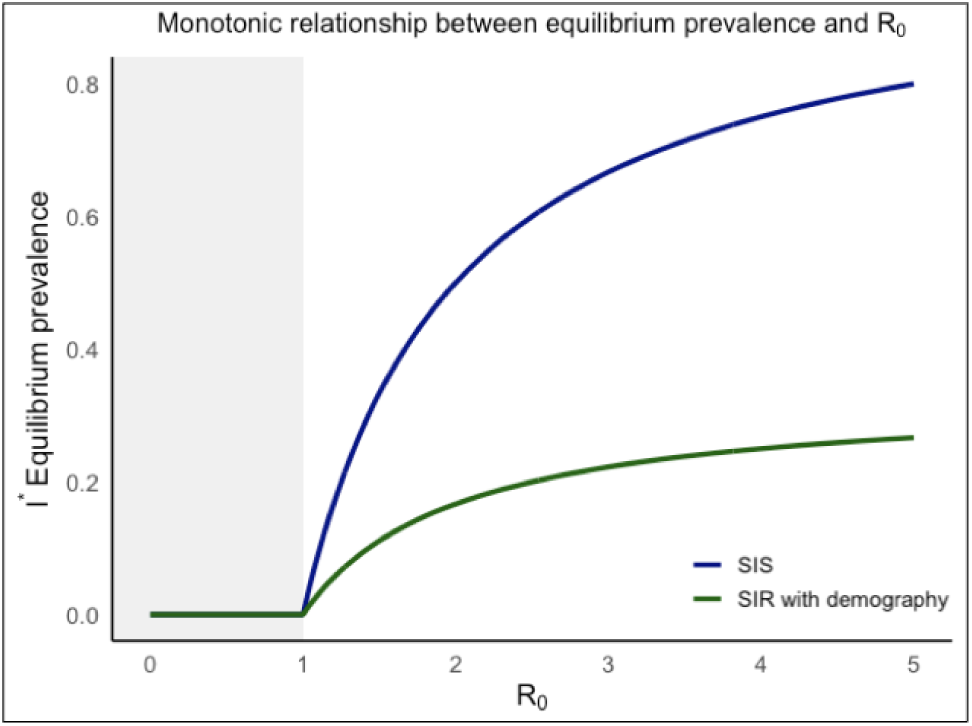

For example, in simple SIR models (Susceptible-Infected-Recovered) with host immunity and demography, equilibrium, prevalence *I*\* satisfies: 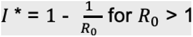.

And by extension to macro parasites:

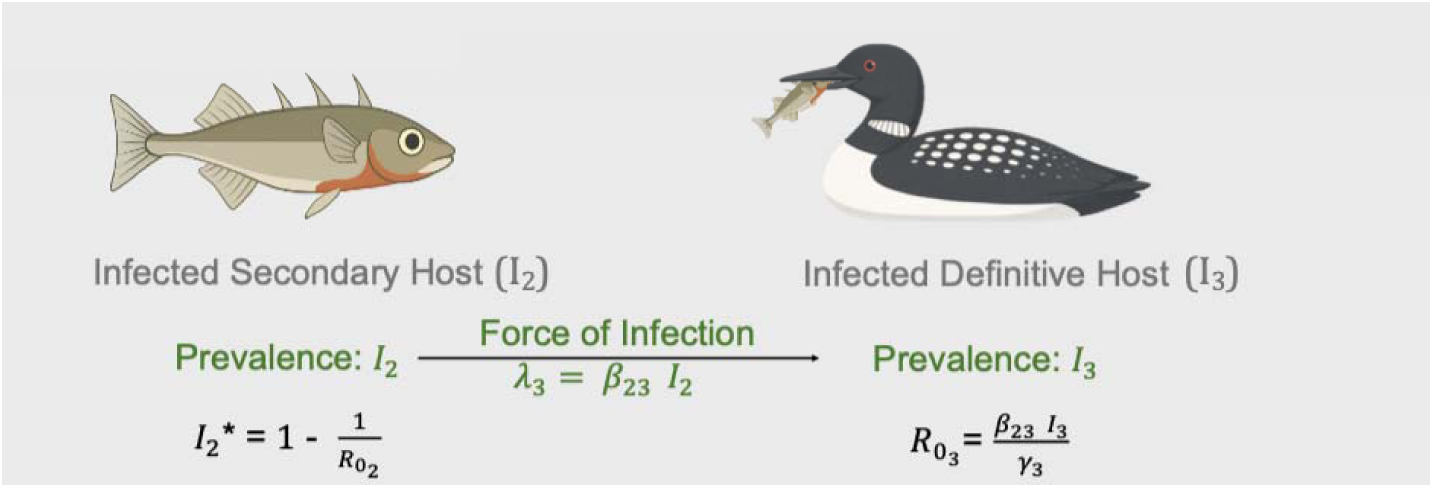

Consequently, all else equal, infection prevalence can capture the same qualitative information as fl_0_, even when transmission rates or *R_o_* are not measured directly due to inherent logistical challenges. Because of this monotonic relationship, a unimodal relationship between virulence and parasite fitness predicted by virulence-transmission trade-off theory will also appear as a unimodal relationship between virulence and infection prevalence. Accordingly, we use infection prevalence as a proxy for parasite fitness to assess whether observed relationships between parasite burden (virulence) and transmission potential to definitive hosts (Fig. **4**) aligns with predictions from virulence-transmission trade-off theory (Alizon et al. 2009, Anderson & May 1982). Rather than challenging the underlying theory, understanding links between empirical estimations of key components of transmission could reveal new opportunities to gain deeper understanding of host-parasite dynamics (McCallum et al. 2017).

