## SUPPLEMENTARY MATERIALS for "Immune variation and host ontogeny shape virulence and transmission potential in a helminth parasite"

Noelle Plantier^4^, Dina Soklovskaya^4^, John Berini III^3^,

Daniel I. Bolnick^4^, Amanda K. Hund^3^, Jessica L. Hite^1^

**Supplementary Table 1.** Monthly stickleback sampling counts by lake.

| **Lake** | **GPS** | **Jul.23** | **Aug** | **Sep** | **Oct** | **Nov** | **Dec** | **Jan** | **Feb** | **Mar** | **Apr** | **May** | **Jun** | **Jul.24** | **Total Fish** |
| --- | --- | --- | --- | --- | --- | --- | --- | --- | --- | --- | --- | --- | --- | --- | --- |
| **BEW** | 48.894931 -124.904225 | 7 | 7 | *NA* | *NA* | *NA* | *NA* | 4 | 3 | 2 | 30 | 23 | 12 | 21 | **109** |
| **BLA** | 48.779385 -125.095096 | 50 | 76 | 96 | 5 | 2 | 10 | 1 | 24 | 60 | 56 | 52 | 52 | 75 | **559** |
| **FRE** | 48.855484 -125.02085 | 50 | 85 | 67 | 13 | 28 | 13 | 2 | 6 | 8 | 55 | 54 | 51 | 60 | **492** |
| **L2B** | 48.813252 -125.127411 | 5 | 3 | 90 | 23 | *NA* | 6 | *NA* | 11 | 7 | 56 | 52 | 52 | 31 | **336** |
| **PCH** | 48.840885 -125.032225 | 52 | 30 | 105 | 71 | 22 | *NA* | *NA* | 3 | 1 | 50 | 51 | 50 | 53 | **488** |
| **RSS** | 48.830346 -125.000446 | 53 | 33 | 28 | 3 | 5 | 5 | 6 | 6 | 13 | 20 | 51 | 50 | 56 | **329** |
| **SRA** | 48.904535 -124.885840 | 101 | 104 | 101 | 17 | 18 | *NA* | 4 | 7 | 8 | 50 | 50 | 52 | 70 | **582** |
| **SUG** | 49.842831 -125.091813 | 69 | 121 | 101 | 8 | 3 | *NA* | *NA* | *NA* | *NA* | 4 | 52 | 49 | 95 | **502** |

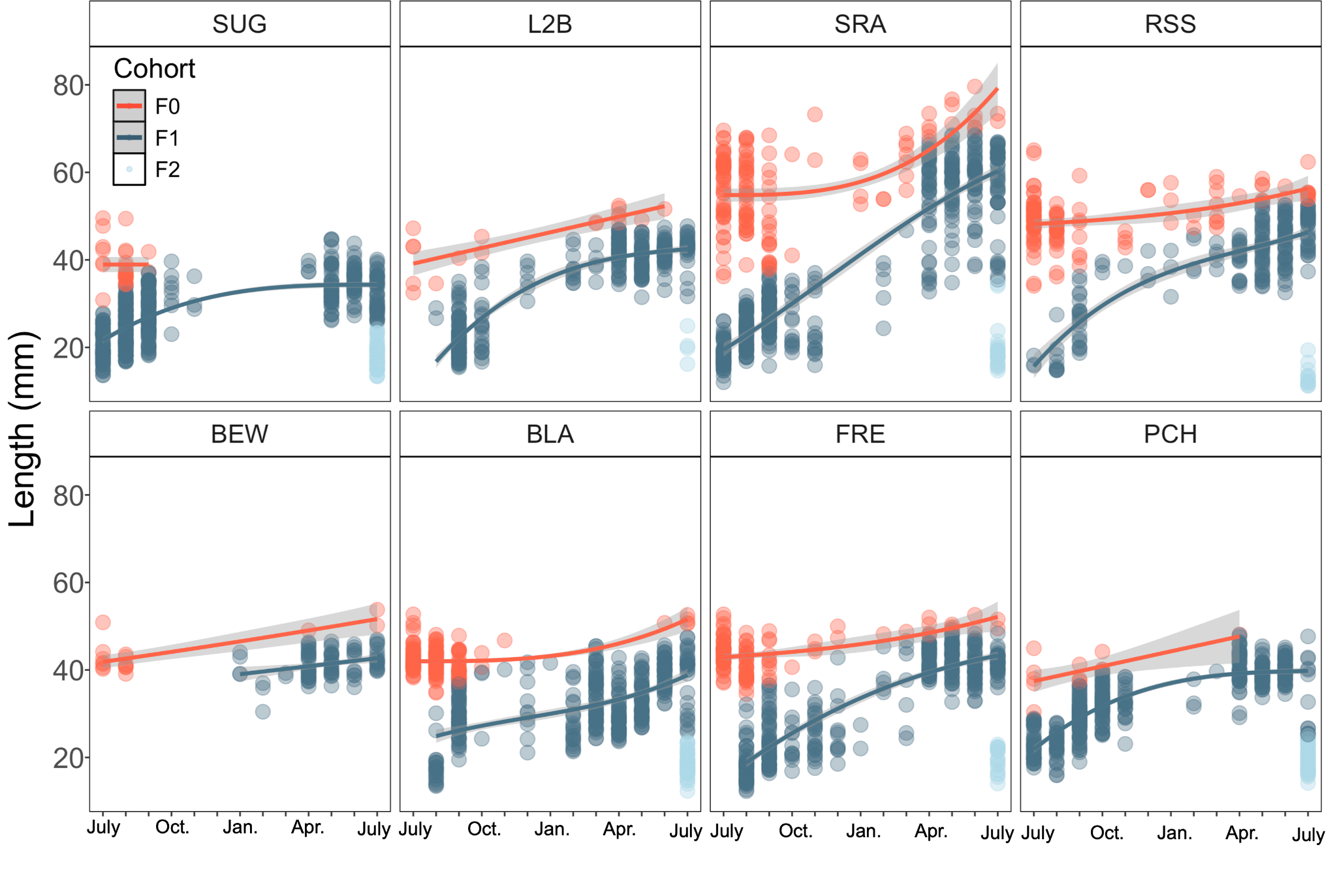

**Supplementary Figure 1. Cohort assignment based on GMM models.**  Each facet represents a lake. Red points denmark F0 generation, dark blue their offspring (F1, the young-of-year that mature across the sampling period) and light blue demarks an emergent F2 population in the following year. Smoothed lines are fit with a monotone increasing smooths fit using a shape-constrained additive model (SCAM).

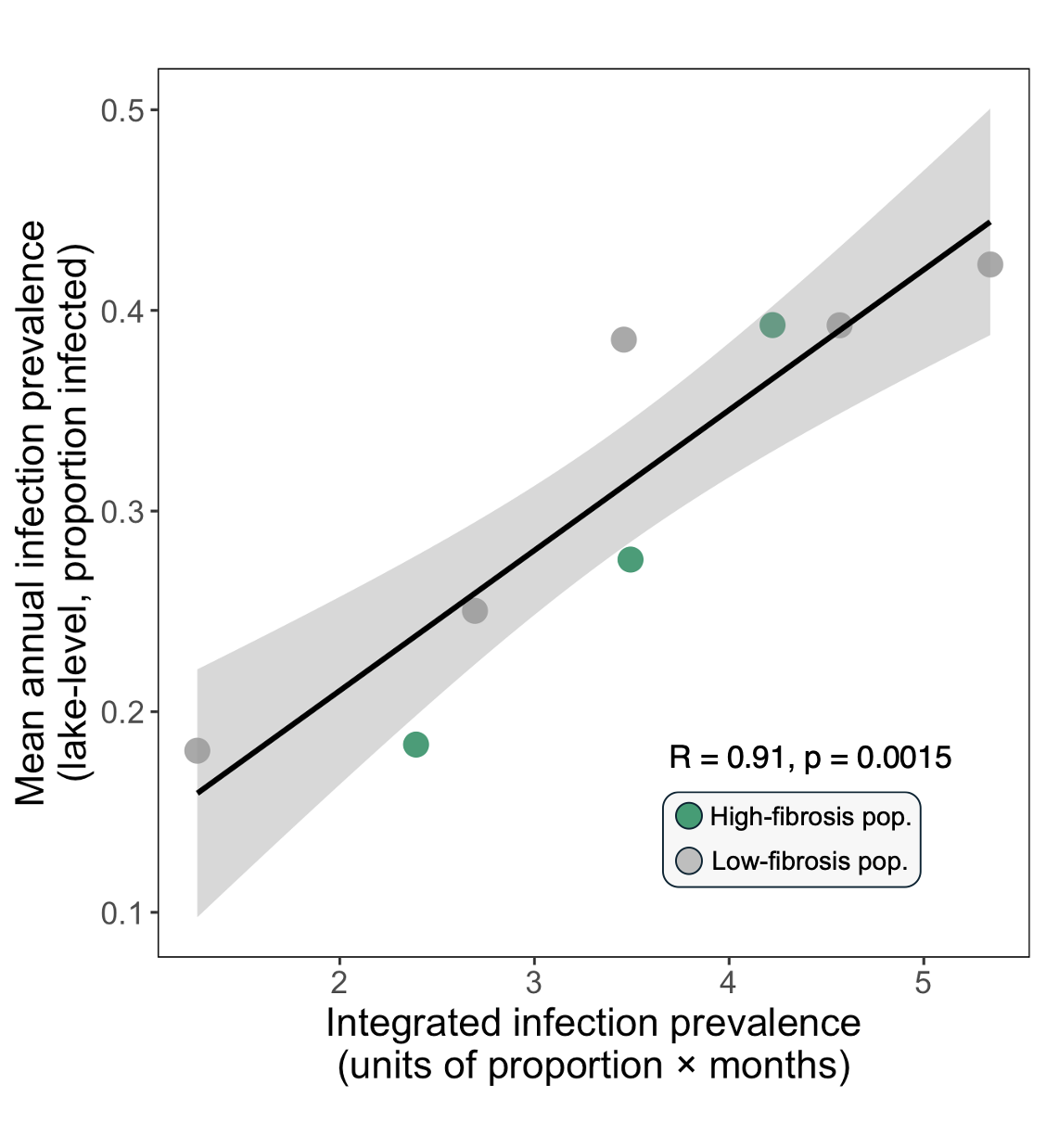

**Supplementary Figure 2. Integrated infection prevalence strongly correlates with maximum infection prevalence.** Points represent lakes, colored by immune phenotype. Smooth line is fit with a glm. Annotation indicates Pearson’s correlation coefficient (R = 0.91, p = 0.0015).

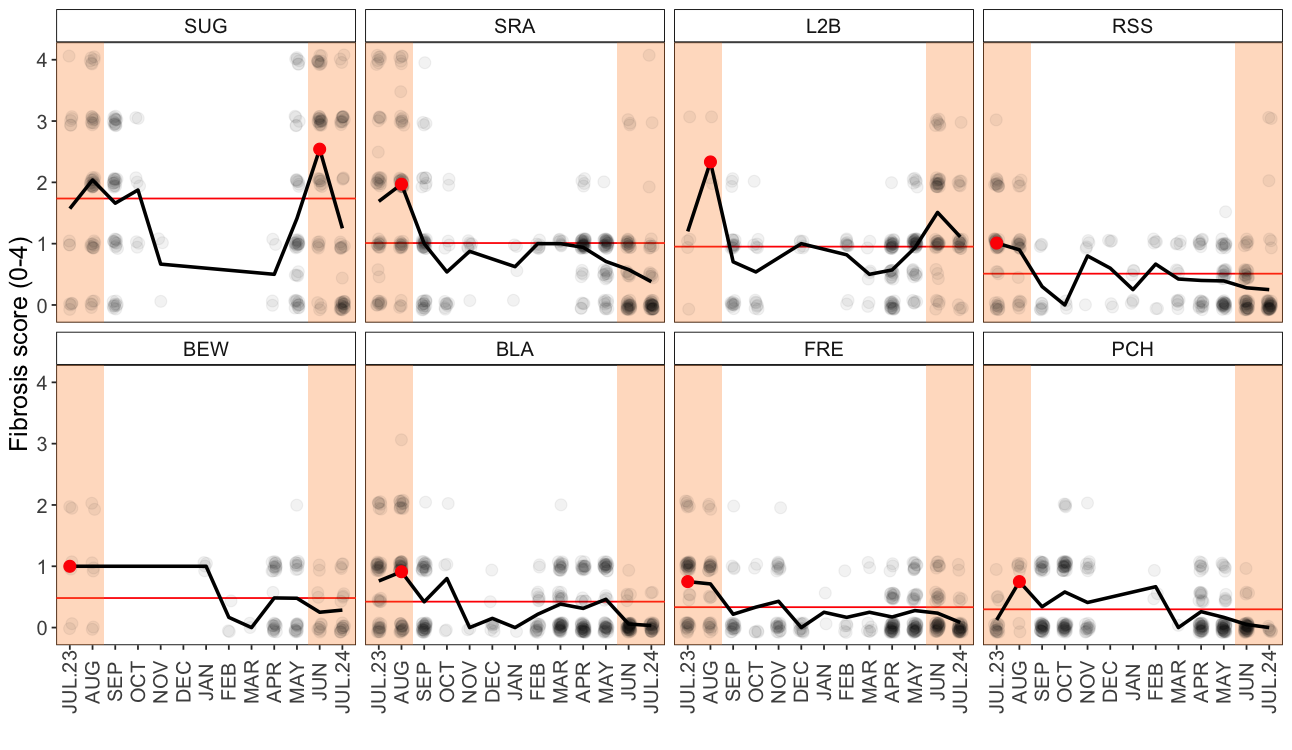

**Supplementary Figure 3. Immune activation peaks in summer months across lakes.** Facets are arranged by decreasing mean fibrosis score. Red lines indicate lake-level annual mean fibrosis. Red points indicate maximum mean fibrosis in each lake across time. Orange panels highlight summer months.

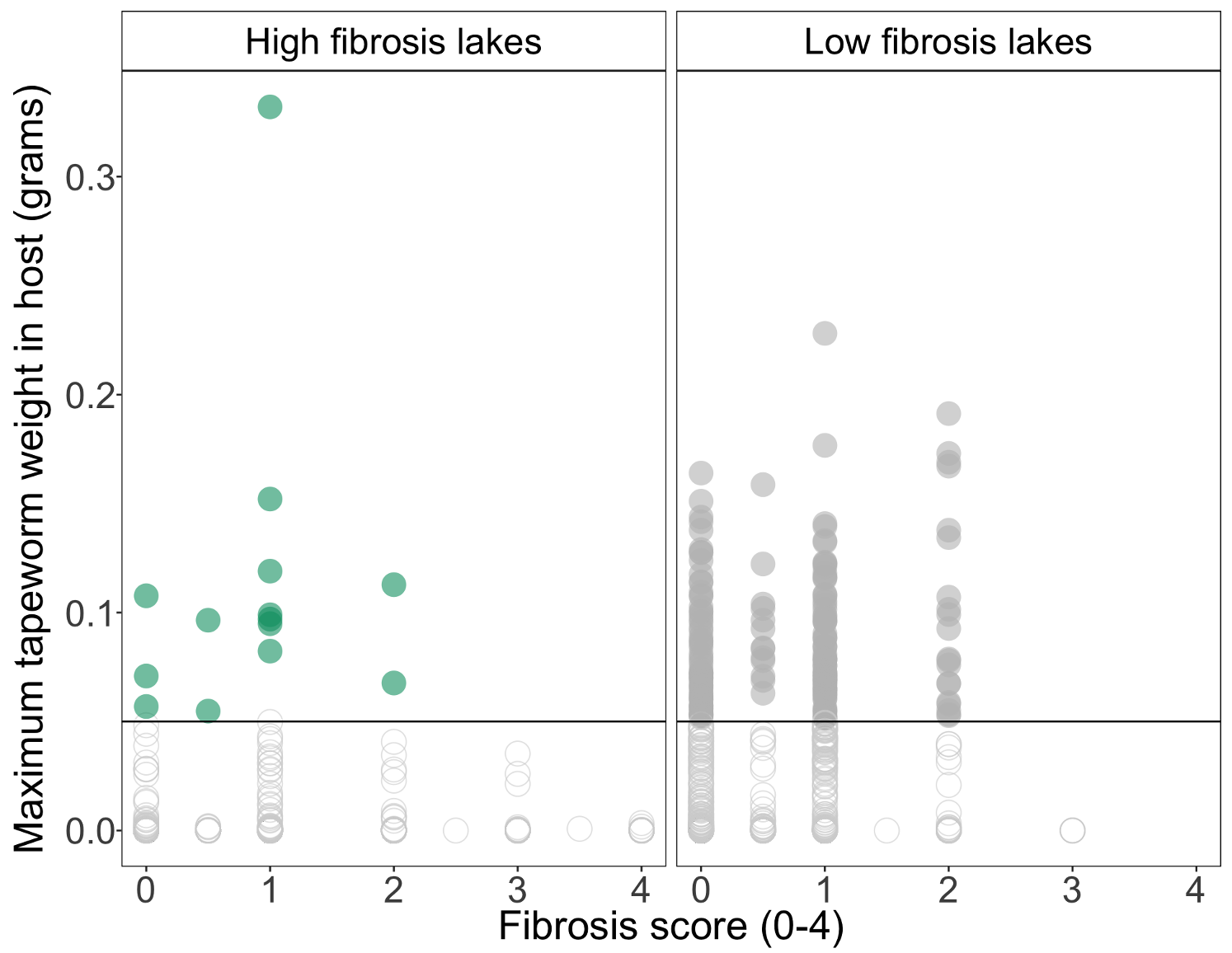

**Supplementary Figure 4.** Sticklebacks that express high fibrosis scores (3 or 4) never contained transmissible parasites. Black line indicates 0.05g threshold at which *S. solidus* can successfully transmit to definitive hosts.

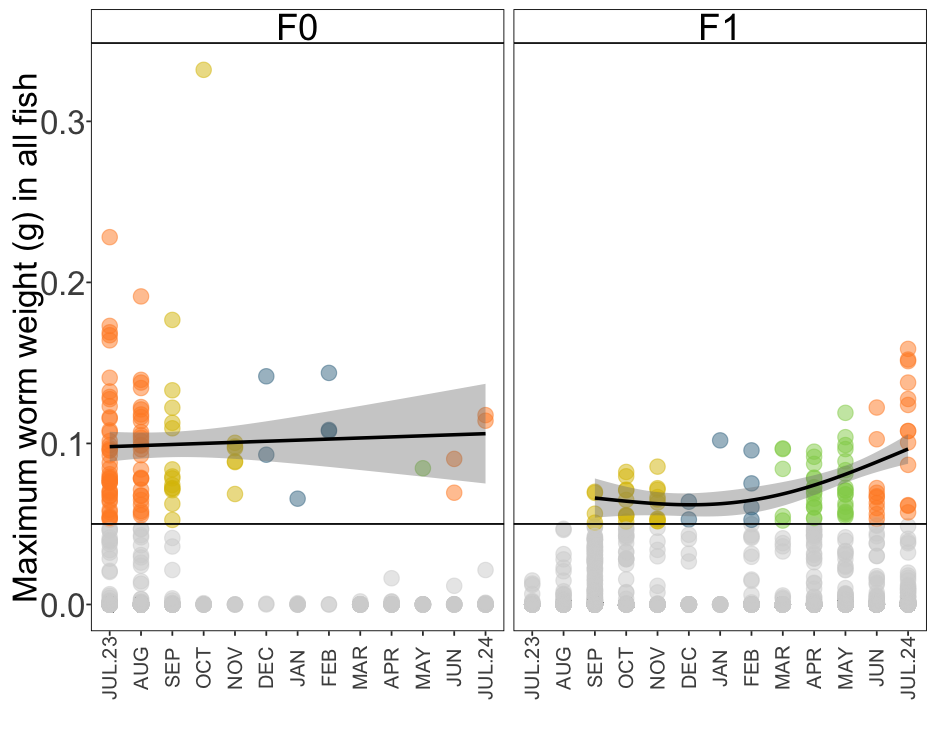

**Supplementary Figure 5.** Transmissible parasite sizes peak in summer months in young-of-year stickleback. Points are colored by season. Black line indicates 0.05g threshold at which tapeworms are transmissible to definitive hosts.
