## Appendix 1 for "Immune variation and host ontogeny shape virulence and transmission potential in a helminth parasite"

**Appendix 1. Phenological analysis of piscovorous bird occurrence**

**Immune variation and host ontogeny constrain virulence and transmission in a helminth parasite.**

Chloe A. Fouilloux^1*^, Heather Alexander^2^, Patrick McNamara^2^, Emma Polard^2^,

Noelle Plantier^4^, Dina Soklovskaya^4^, John Berini III^3^,

Daniel I. Bolnick^4^, Amanda K. Hund^3^, Jessica L. Hite^1^

**Field Collection and Sampling Design**

We deployed autonomous recording units (ARU; Wildlife Acoustics Song Meter SM4) at eight lake sites in the Bamfield area of Vancouver Island, British Columbia. ARUs were mounted on vertical structures at heights of 1 to 3 meters. To capture a comprehensive record of the avian soundscape, ARUs were programmed on a duty cycle of 30 minutes on and 30 minutes off, 24 hours a day, from August 1, 2023, to July 31, 2024. To maximize storage capacity during these extended deployments, we utilized Wildlife Acoustics’ proprietary W4V-8 compression option, which doubled the potential recording length while maintaining a high signal-to-noise ratio. To ensure data integrity and facilitate site identification, field personnel provided verbal metadata, including name, date, time, and GPS coordinates, directly into the microphone at the initiation and conclusion of every recording session, complementing the standard file-naming conventions established during unit deployment. Audio recordings were processed using BirdNET (Wood et al. 2022), a software that detects and classifies avian sounds using machine learning. Batteries were replaced and SD memory card data were downloaded monthly.

**Data Decompression and Conversion**

Because we chose to record audio in a proprietary compressed format, raw files were processed using Wildlife Acoustics’ Kaleidoscope Lite Version 5.6.8 (Wildlife Acoustics, Inc. 2026) in "Non-bat Analysis Mode." We utilized the batch processing function to decompress the .w4v files into standard .wav format. To ensure no data loss or signal degradation during conversion, we configured the software to “process all input channels” and set the output compression to "None," providing the uncompressed signal necessary for high-accuracy automated identification.

**Automated Species Identification (BirdNET)**

To process the more than 10,000 hours of audio collected across our eight sites, we utilized the BirdNET-Analyzer GUI v2.4 (Kahl et al. 2021), a deep-learning convolutional neural network developed by the Cornell Lab of Ornithology and Chemnitz University of Technology. The algorithm partitioned the raw audio into consecutive three-second segments, converting them into visual spectrograms for identification.

To optimize identification accuracy and mitigate the risk of false positives, we implemented two specific BirdNET constraints – Species by Location Filtering and Confidence Thresholding.

**Species-by-Location Filtering:** We utilized the "Species by Location" feature, entering coordinates for the Bamfield region (Lat: 48.843336, Lon: -125.018680) and a temporal filter centered on the 24th week of the year (June) to align the model’s internal probability with the known phenology of the study area.

**Confidence Thresholding:** We applied a minimum confidence threshold of 0.75. This value was selected based on meta-analytical findings (e.g., Perez-Granados, 2023) suggesting that 0.75 provides the optimal balance between recall and precision across diverse habitat types.

Because BirdNET generates a unique detection log for every audio file processed (in our case, a discrete .csv for every 30-minute recording), this stage resulted in thousands of files. To handle this volume, we utilized parallel processing, adjusting batch sizes and threads to match the capabilities of the specific multicore processor being employed. It is important to note that these data represent acoustic activity rather than absolute abundance; because ARUs cannot distinguish between multiple calls made by a single individual and single calls made by multiple individuals, our metrics reflect the frequency of detection.

**Data Integration and Metadata Reconstruction**

The fragmented output from BirdNET lacked inherent site-level metadata within the individual data rows. We developed a custom R script to "mine" the necessary variables from the directory structure and file naming conventions. This script parsed the file path strings to extract and assign variables for the monitor ID, location code, and recording date, merging thousands of individual one half-hour-long logs into a unified, site-specific database.

**Temporal Processing and Precision Timing**

We leveraged the intrinsic temporal logging capabilities of the SM4 to ensure high-resolution phenological analysis. The devices automatically embedded start-time metadata into each file header. By programmatically adding the BirdNET detection offset (in seconds) to these baseline timestamps, we assigned a discrete, real-world timestamp to every vocalization (e.g., a detection at a 33-second offset in a 09:00:00 recording was logged as 09:00:33). We validated the continuity of this effort through a systematic gap analysis, mapping actual versus expected recording frequency to ensure that "zeros" in the dataset represented biological silence rather than hardware downtime.

**Longitudinal Aggregation and Matrix Transformation**

For the final processing stage, we summarized the detection rows into weekly and monthly counts. Using R, we reshaped these data into a longitudinal matrix, where each row represents a unique site-week combination and each column represents a specific bird species. During this transformation, we standardized site nomenclature (e.g., correcting mismatched site identifiers) to ensure the resulting matrix was appropriately formatted for subsequent ecological modeling and biodiversity analysis.

**Sources**

Wildlife Acoustics, Inc. 2026. Kaleidoscope Lite (Version 5.6.8) [Computer software]. https://www.wildlifeacoustics.com/products/kaleidoscope/kaleidoscope-lite

Kahl, S., Wood, C. M., Eibl, M. and H. Klinck. 2021. BirdNET: A deep learning solution for avian diversity monitoring. Ecological Informatics 61: 101236.

Perez-Granados, C. 2023. BirdNET: applications, performance, pitfalls and future opportunities. IBIS 165(3): 1068-1075.

Github repo: <https://github.com/jberini/BC_ARU_longitudinal_processing>

Contains:

- README.md
- Script
- Sample data
