## Appendix 2 for "Immune variation and host ontogeny shape virulence and transmission potential in a helminth parasite"

**APPENDIX 2. Detailed Statistical Methods**

**Immune variation and host ontogeny constrain virulence and transmission in a helminth parasite**

Chloe A. Fouilloux^1*^, Heather Alexander^2^, Patrick McNamara^2^, Emma Polard^2^,

Noelle Plantier^4^, Dina Soklovskaya^4^, John Berini III^3^,

Daniel I. Bolnick^4^, Amanda K. Hund^3^, Jessica L. Hite^1^

**Detailed statistical analyses.**

***Identifying immune phenotypes across host populations –*** To contextualize parasite dynamics within their host populations, we classified lakes by immune phenotype using partitioning around medoids (PAM). Because fibrosis reflects an induced immune response to parasite exposure, classification integrated both host immune data and infection data. For each lake, we calculated mean fibrosis scores, infection prevalence (proportion of fish harboring ≥1 tapeworm), and mean parasite burden (total mass of tapeworms (g) divided by the mass of the host (g)) (Table 1). We constructed a Gower dissimilarity matrix using daisy (cluster v2.1.8.1, Maechler et al. 2026) with variable-weighted dissimilarity, and computed cluster separation across k = 2 to 5; k = 2 maximised separation and was therefore selected. These dissimilarities were passed to the PAM algorithm (pam), yielding two discrete lake groups: high- versus low-fibrosis populations (Table 1). These population-level summaries, henceforth called immune phenotypes, were aggregated across all sampling periods and treated as fixed group labels in downstream models. For each lake, we calculated mean fibrosis scores, infection prevalence (proportion of fish harboring ≥1 tapeworm), and mean parasite burden (Table 1).

***Identifying cohorts in naturally occurring populations –*** Immune defense and parasite growth both shift across host ontogeny, so characterising population-level patterns required classifying sampled fish into generational cohorts. To do so, we classified fish into generational cohorts using a model‑based clustering approach informed by size‑at‑age dynamics. To ensure biological relevance, cohort assignments were constrained using month-specific size caps derived from growth expectations based on dynamic energy budget theory growth expectations (Wright et al. 2004). For lakes with atypically large fish (SRA, RSS; see [Supp. Fig 1](https://docs.google.com/document/d/1HXIHI3oQm34lUHHfh9ZeFH7J5G6jLCVBljAn4r5_6Po/edit?tab=t.0) for lake-specific cohort size distributions), month-specific size caps were increased by a fixed offset to reflect local growth conditions.

Host populations differed markedly in body size distributions (Kruskal-Wallis test, X2 = 553.3, p < 0.001), reflecting variation in productivity and growth environments. To capture this heterogeneity, we classified fish within each lake and month based on length distributions. Initial cohort structure was identified using early-season data (July-September 2023), when only two cohorts are expected (F0 adults and young-of-year F1), assuming an annual generation cycle. For each lake-month combination, we fit Gaussian mixture models (Mclust, v6.1.1, Scrucca et al. 2023) with up to two components, allowing variance to differ among components. Components were ranked by mean length and labeled as F1 (smaller) or F0 (larger); in months 12-13 (June-July 2024) we allowed up to three components to capture the expected emergence of F2 recruits. Mixture components were again ordered by mean length, and provisional cohort labels were assigned by matching each fish to the nearest monthly component mean. In lake-month combinations with insufficient data for mixture modelling, we used k-means clustering, which assigns each fish to the nearest estimated cohort mean without modelling variance or mixing weights. F2 assignments were restricted to individuals below a fixed size threshold in months 12-13; fish exceeding the monthly F1 size cap were reassigned to F0, and fish falling below it were reassigned to F1.

Although F2 cohorts were detectable in some lakes by July 2024, our analyses focus on F0 and F1 cohorts, which encompass the majority of infection and immune data; F2 fish (~5.5% of sampled fish) were excluded from all analyses. All steps were applied independently to each lake and month, allowing cohort boundaries to shift with local growth conditions

***Q1. How does the timing and magnitude of parasite burden (a proxy for virulence) vary across seasons and among host populations that differ in immune phenotype? –***

Using the stage structure and immune phenotype classifications outlined above, we fit a Bayesian zero-inflated beta regression (brms v2.23.0, Bürkner 2021; RStan v2.32.7) to model within-host parasite burden across host populations and through time. To capture nonlinear seasonal trajectories independently for each immune phenotype, the model included three submodels: (i) the mean submodel (mu) included immune phenotype, host cohort, and immune-phenotype-specific thin-plate regression splines across time; (ii) the zero-inflation (zi) and precision (phi) submodels each included immune phenotype, a lake-level random intercept, and immune-phenotype-specific thin-plate regression splines across time. Lake was excluded from the mean submodel because lakes were perfectly nested within immune phenotype, precluding simultaneous estimation of lake-level random effects and population-level fibrosis differences in the mean structure.

Weakly informative priors were specified for all parameters: student-t(3, 0, 2.5) for intercepts, normal(0, 0.6) for fixed effects, exponential(1) for smoothing standard deviations, normal(log(40), 0.6) for the phi intercept (reflecting prior expectation of moderate precision based on the bounded nature of the response), and student-t(3, 0, 2.5) for random effect standard deviations in the zi and phi submodels. Four MCMC chains were run for 3000 iterations each with 1000 warmup iterations (adapt_delta = 0.99, max_treedepth = 14), yielding 8000 post-warmup draws.

Model convergence was assessed via the potential scale reduction factor (all Rhat ≤ 1.01) and effective sample size ratios (bulk ESS > 1000 for all parameters). Model adequacy was evaluated using DHARMa-style posterior predictive simulations (DHARMa v0.4.7, Hartig 2024), which indicated acceptable fit (Kolmogorov-Smirnov test nonsignificant, no outliers). A marginal degree of underdispersion was detected (dispersion ratio = 0.71, p = 0.048), attributable to the highly zero-skewed burden distribution in high-fibrosis populations. No residual temporal autocorrelation based on Moran’s I (p = 0.06). Leave-one-out cross-validation confirmed adequate predictive performance with all Pareto k estimates below 0.7.

***Q2. How does the timing and magnitude of host immune response vary across seasons, host ontogeny and immune phenotype? –***  We assessed how immune expression varied as a function of immune phenotype, host cohort, and infection status across time. Individual fibrosis was analyzed using a generalized additive mixed model implemented in the mgcv package (v. 1.9.3, Wood et al. 2011). Models were fitted using restricted maximum likelihood (REML). Model diagnostics using DHARMa indicated no issues with dispersion, residual structure, or outliers. Of note, compared to summer months, winter months were sparsely sampled ([Supp Table 1](https://docs.google.com/document/d/1HXIHI3oQm34lUHHfh9ZeFH7J5G6jLCVBljAn4r5_6Po/edit?tab=t.0)). As a result, uncertainty intervals in November-March are wider and should be interpreted with caution.

***Q3. Do immune‑mediated differences in parasite burden scale up to influence transmission potential to definitive hosts? –*** We examined whether immune-phenotype-driven differences in transmissible parasite production align with peaks in avian definitive host richness, linking within-host immune dynamics to population-level transmission potential.

Spearman rank correlations were used to test the association between monthly piscivorous bird species richness and transmissible parasite counts, computed separately for high- vs. low-fibrosis populations (n = 13 months per group). Because monthly transmissible parasite counts differed substantially in absolute magnitude between immune populations (range: 0-42), counts were standardized (z-score: (x−μ)/σ) within each immune phenotype prior to visualization to facilitate comparison of seasonal timing and correlation structure independently of differences in magnitude (Fig. 3H).
